## Supplemental Info for "A portable regulatory RNA array design enables tunable and complex regulation across diverse bacteria"

### Table of contents


| Tables | Title | Page |
|---|---|---|
| 1 | All plasmids used in this study | 2 - 5 |
| 2 | Example DNA plasmid sequence | 6 – 10 |
| 3 | SP6 promoter sequences used in this study | 11 |
| 4 | Golden Gate assembly primers and overhangs | 12 |
| 5 | Strains, growth conditions, and electroporation settings used in this study | 13 |
| 6 | Regulatory RNA array insulator sequences | 14 |
| **Figures** | | |
| 1 | Characterization of a STAR system that uses the SP6 RNA polymerase | 15 |
| 2 | A modular cloning (MoClo) approach for regulatory RNA arrays | 16 |
| 3 | Insulator screening for regulatory RNA arrays | 17 |
| 4 | NUPACK analysis of regulatory RNA arrays with different insulators | 18 |
| 5 | Calibration curves of inducible promoters to relate promoter strength to transcriptional output in different bacterial species | 19 |
| 6 | The induction curves of the regulatory RNA array before the promoter activity normalization | 20 |
| 7 | The schematic of an RNA activation-activation cascade | 21 |
| 8 | Fluorescent characterization of multiplex RNA arrays | 22 |
| 9 | Schematic of representative plasmids used in this study | 23 |
| **Notes** | | |
| 1 | Mathematical model of regulatory RNA arrays | 24 - 25 |

**Supplementary Table 1. All plasmids used in this study.** CmR: chloramphenicol resistance gene, AmpR: ampicillin resistance gene, SpcR: spectinomycin resistance gene, KanR: kanamycin resistance gene, p15A: origin of replication, ColE1: origin of replication, CDF: CloDF origin of replication, pBBR1: broad-host range origin of replication, pSP6: SP6 promoter, TrrnB: terminator, t500: terminator, RBS: ribosome binding site, Csy4: Csy4 ribonuclease, csy4hp: csy4 cleavage hairpin, shcsyhp: strong hairpin csy4 cleavage hairpin, PlmJ: ribozyme.

| Plasmid | Description | Plasmid Architect | Figure |
|---|---|---|---|
| pBL176 | STAR positive | J23119-Target10-RBS-GFP-TrrnB-J23119-STAR10-t500-pBBR1-KanR | Fig1 |
| pBL177 | STAR negative | J23119-Target10-RBS-GFP-TrrnB-pBBR1-KanR | Fig1 |
| pBL301 | STAR negative+Csy4 | J23119-Target10-RBS-GFP-TrrnB-t500-J23101-Csy4-Terminator-pBBR1-KanR | Fig1/Fig4 |
| pBL302 | STAR positive+Csy4 | J23119-Target10-RBS-GFP-TrrnB-J23119-STAR10-t500-J23101-Csy4-Terminator-pBBR1-KanR | Fig1 |
| pBL115 | SP6 RNAP | AraC-pBAD-RBS-SP6 RNAP-TrrnB-SpcR-CDF | Fig1/FigS1 |
| pBL068 | SP6 Target-GFP | pSP6-Target6-RBS-GFP-TrrnB-CmR-p15A | Fig1 |
| pBL071 | SP6 STAR | pSP6-STAR6-t500-ColE1-AmpR | Fig1 |
| pJEC102 | Control | J23119-TrrnB-ColE1-AmpR | Fig1/FigS1 |
| pJEC250 | Target50-GFP | J23119-Target50-RBS-GFP-TrrnB-CmR-p15A | Fig2/FigS3 |
| pBLM443 | PlmJ_STAR50x4 | AraC_pBAD-PlmJ_STAR50x4-t500-ColE1-AmpR | Fig2 |
| pBLM444 | PlmJ_STAR50x1 | AraC_pBAD-PlmJ_STAR50x1-t500-ColE1-AmpR | Fig2 |
| pBL111 | Csy4 | J23101-Csy4-CDF-SpcR | Fig2 |
| pBL451 | csy4hp_STAR50x4 | J23119-Target50-RBS-GFP-TrrnB-AraC_pBAD-csy4hp_STAR50x4-CmR-p15A | Fig2 |
| pBL452 | csy4hp_STAR50x1 | J23119-Target50-RBS-GFP-TrrnB-AraC_pBAD-csy4hp_STAR50x1-CmR-p15A | Fig2 |
| pBL170 | shcsy4hp_STAR50x4 | J23119-Target50-RBS-GFP-TrrnB-AraC_pBAD-shcsy4hp_STAR50x4-CmR-p15A | Fig2 |
| pBL171 | shcsy4hp_STAR50x1 | J23119-Target50-RBS-GFP-TrrnB-AraC_pBAD-shcsy4hp_STAR50x1-CmR-p15A | Fig2 |

| | | | |
|---|---|---|---|
| pBL161 | shcsy4hp_STAR10x4 | J23119-Target10-RBS-GFP-TrrnB-AraC_pBAD-shcsy4hp_STAR10x4-CmR-p15A | Fig2 |
| pBL162 | shcsy4hp_STAR10x1 | J23119-Target10-RBS-GFP-TrrnB-AraC_pBAD-shcsy4hp_STAR10x1-CmR-p15A | Fig2 |
| pJEC216 | Target10-GFP | J23119-Target10-RBS-GFP-TrrnB-CmR-p15A | Fig2/Fig4/FigS3 |
| pBL303 | STAR10x1 (pBAD) | J23119-Target10-RBS-GFP-TrrnB-AraC_pBAD-shcsy4hp_STAR10x1-J23101-Csy4-Terminator-pBBR1-KanR | Fig3/FigS6 |
| pBL304 | STAR10x2 (pBAD) | J23119-Target10-RBS-GFP-TrrnB-AraC_pBAD-shcsy4hp_STAR10x2-J23101-Csy4-Terminator-pBBR1-KanR | Fig3/FigS6 |
| pBL305 | STAR10x4 (pBAD) | J23119-Target10-RBS-GFP-TrrnB-AraC_pBAD-shcsy4hp_STAR10x4-J23101-Csy4-Terminator-pBBR1-KanR | Fig3/FigS6 |
| pBL306 | STAR10x6 (pBAD) | J23119-Target10-RBS-GFP-TrrnB-AraC_pBAD-shcsy4hp_STAR10x6-J23101-Csy4-Terminator-pBBR1-KanR | Fig3/FigS6 |
| pBL307 | STAR10x8 (pBAD) | J23119-Target10-RBS-GFP-TrrnB-AraC_pBAD-shcsy4hp_STAR10x8-J23101-Csy4-Terminator-pBBR1-KanR | Fig3/FigS6 |
| pBL401 | STAR10x1 (pCymR) | J23119-Target10-RBS-GFP-TrrnB-CymR_pCymR-shcsy4hp_STAR10x1-J23101-Csy4-Terminator-pBBR1-KanR | Fig3/FigS6 |
| pBL402 | STAR10x2 (pCymR) | J23119-Target10-RBS-GFP-TrrnB-CymR_pCymR-shcsy4hp_STAR10x2-J23101-Csy4-Terminator-pBBR1-KanR | Fig3/FigS6 |
| pBL403 | STAR10x4 (pCymR) | J23119-Target10-RBS-GFP-TrrnB-CymR_pCymR-shcsy4hp_STAR10x4-J23101-Csy4-Terminator-pBBR1-KanR | Fig3/FigS6 |
| pBL404 | STAR10x6 (pCymR) | J23119-Target10-RBS-GFP-TrrnB-CymR_pCymR-shcsy4hp_STAR10x6-J23101-Csy4-Terminator-pBBR1-KanR | Fig3/FigS6 |
| pBL405 | STAR10x8 (pCymR) | J23119-Target10-RBS-GFP-TrrnB-CymR_pCymR-shcsy4hp_STAR10x8-J23101-Csy4-Terminator-pBBR1-KanR | Fig3/FigS6 |
| pBL512 | pBAD-GFP | AraC_pBAD-RBS-GFP-TrrnB-pBBR1-KanR | Fig3/FigS5 |

| | | | |
|---|---|---|---|
| pBL513 | pCymR-GFP | CymR_pCymR-RBS-GFP-TrrnB-pBBR1-KanR | Fig3/FigS5 |
| pBLM366 | Target50-STAR10x1 | J23119-Target50-shCsy4_STAR10x1-t500-RSF1030-SpcR | Fig4 |
| pBLM367 | Target50-STAR10x4 | J23119-Target50-shCsy4_STAR10x4-t500-RSF1030-SpcR | Fig4 |
| pJEC251 | STAR50 | J23119-STAR50-t500-ColE1-AmpR | Fig4 |
| pBL501 | Target10-GFP+Target50-STAR10x1 | J23119-Target10-RBS-GFP-TrrnB-J23119-Target50-shCsy4_STAR10x1-t500-J23101-Csy4-Terminator-pBBR1-KanR | Fig4 |
| pBL502 | Target10-GFP+Target50-STAR10x4 | J23119-Target10-RBS-GFP-TrrnB-J23119-Target50-shCsy4_STAR10x4-t500-J23101-Csy4-Terminator-pBBR1-KanR | Fig4 |
| pBL504 | Target10-GFP+Target50-STAR10x1 +STAR50 | J23119-Target10-RBS-GFP-TrrnB-J23119-Target50-shCsy4_STAR10x1-t500-J23119-STAR50-t500-J23101-Csy4-Terminator-pBBR1-KanR | Fig4 |
| pBL505 | Target10-GFP+Target50-STAR10x4 +STAR50 | J23119-Target10-RBS-GFP-TrrnB-J23119-Target50-shCsy4_STAR10x4-t500-J23119-STAR50-t500-J23101-Csy4-Terminator-pBBR1-KanR | Fig4 |
| pBL506 | Target10-GFP+Target50-mRFP+STAR10x1 +STAR50x1 | J23119-Target10-RBS-GFP-TrrnB-J23119-RBS-RFP-t500-AraC_pBAD-shcsy4hp_STAR10x1-shcsy4hp_STAR50x1-t500-J23101-Csy4-Terminator-pBBR1-KanR | Fig4/FigS8 |
| pBL507 | Target10-GFP+Target50-mRFP+STAR10x1 +STAR50x2 | J23119-Target10-RBS-GFP-TrrnB-J23119-RBS-RFP-t500-AraC_pBAD-shcsy4hp_STAR10x1-shcsy4hp_STAR50x2-t500-J23101-Csy4-Terminator-pBBR1-KanR | Fig4/FigS8 |
| pBL508 | Target10-GFP+Target50-mRFP+STAR10x2 +STAR50x1 | J23119-Target10-RBS-GFP-TrrnB-J23119-RBS-RFP-t500-AraC_pBAD-shcsy4hp_STAR10x2-shcsy4hp_STAR50x1-t500-J23101-Csy4-Terminator-pBBR1-KanR | Fig4/FigS8 |
| pBL509 | Target10-GFP+Target50-mRFP+STAR10x2 +STAR50x2 | J23119-Target10-RBS-GFP-TrrnB-J23119-RBS-RFP-t500-AraC_pBAD-shcsy4hp_STAR10x2-shcsy4hp_STAR50x2-t500-J23101-Csy4-Terminator-pBBR1-KanR | Fig4/FigS8 |
| pBL059 | SP6 RNAP | pTet-RBS-luxR-Terminator-pLuxR-RBS-SP6 RNAP-TrrnB-SpcR-CDF | FigS1 |

| | | | |
|---|---|---|---|
| pBL062 | SP6 promoter-Target28-GFP | pSP6-Target28(AD1.S7)-RBS-sfGFP-TrrnB-CmR-p15A | FigS1 |
| pJEC154 | STAR28(12nt) | J23119-STAR28(12nt)-t500-ColE1-AmpR | FigS1 |
| pJEC163 | STAR28(17nt) | J23119-STAR28(17nt)-t500-ColE1-AmpR | FigS1 |
| pJEC164 | STAR28(27nt) | J23119-STAR28(27nt)-t500-ColE1-AmpR | FigS1 |
| pJEC165 | STAR28(37nt) | J23119-STAR28(37nt)-t500-ColE1-AmpR | FigS1 |
| pJEC166 | STAR28(47nt) | J23119-STAR28(47nt)-t500-ColE1-AmpR | FigS1 |
| pJEC167 | STAR28(57nt) | J23119-STAR28(57nt)-t500-ColE1-AmpR | FigS1 |
| pBL116 | pSP6(wt)-Target10-GFP | pSP6-Target10-RBS-GFP-TrrnB-CmR-p15A | FigS1 |
| pBL120 | pSP6(wt)-STAR10 | pSP6-STAR10-t500-ColE1-AmpR | FigS1 |
| pBL144 | pSP6(90p)-STAR10 | pSP6(90p)-STAR10-t500-ColE1-AmpR | FigS1 |
| pBL145 | pSP6(50p)-STAR10 | pSP6(50p)-STAR10-t500-ColE1-AmpR | FigS1 |
| pBL146 | pSP6(20p)-STAR10 | pSP6(20p)-STAR10-t500-ColE1-AmpR | FigS1 |
| pBL117 | pSP6-Target28-GFP | pSP6-Target28-RBS-GFP-TrrnB-CmR-p15A | FigS1 |
| pBL118 | pSP6-Target50-GFP | pSP6-Target50-RBS-GFP-TrrnB-CmR-p15A | FigS1 |
| pBL121 | pSP6-STAR28 | pSP6-STAR28-t500-ColE1-AmpR | FigS1 |
| pBL122 | pSP6-STAR50 | pSP6-STAR50-t500-ColE1-AmpR | FigS1 |

**Supplementary Table 2. Example DNA plasmid sequence.**

| Name | DNA sequence |
|---|---|
| Regulatory RNA array plasmid STAR10x8 (pBL307) J23119-Target10-RBS-GFP-TrrnB-AraC-pBAD-(shcsy4hp_STAR10)x8-Csy4-pBBR1-KanR | GAATTCTAAAGATCTTTGACAGCTAGCTCAGTCCTAGGTATAATACTAGT<br>CTCATCTCATTTCGCTCTCACATTTCTACACCTTTATCTTGCGGGGAATGT<br>ATACAGTTCATGTATATATTCCCCGCTTTTTTTTTGGATCTAGGAGGAAG<br>GATCTATGAGCAAAGGAGAAGAACTTTTCACTGGAGTTGTCCCAATTCTT<br>GTTGAATTAGATGGTGATGTTAATGGGCACAAATTTTCTGTCCGTGGAGA<br>GGGTGAAGGTGATGCTACAAACGGAAAACTCACCCTTAAATTTATTTGCA<br>CTACTGGAAAACTACCTGTTCCGTGGCCAACACTTGTCACTACTCTGACC<br>TATGGTGTTCAATGCTTTTCCCGTTATCCGGATCACATGAAACGGCATGA<br>CTTTTTCAAGAGTGCCATGCCCGAAGGTTATGTACAGGAACGCACTATAT<br>CTTTCAAAGATGACGGGACCTACAAGACGCGTGCTGAAGTCAAGTTTGA<br>AGGTGATACCCTTGTTAATCGTATCGAGTTAAAGGGTATTGATTTTAAAG<br>AAGATGGAAACATTCTTGGACACAAACTCGAGTACAACTTTAACTCACAC<br>AATGTATACATCACGGCAGACAAACAAAAGAATGGAATCAAAGCTAACTT<br>CAAAATTCGCCACAACGTTGAAGATGGTTCCGTTCAACTAGCAGACCATT<br>ATCAACAAAATACTCCAATTGGCGATGGCCCTGTCCTTTTACCAGACAAC<br>CATTACCTGTCGACACAATCTGTCCTTTCGAAAGATCCCAACGAAAAGCG<br>TGACCACATGGTCCTTCTTGAGTTTGTAACTGCTGCTGGGATTACACATG<br>GCATGGATGAGCTCTACAAATAAGGATCTGAAGCTTGGGCCCGAACAAA<br>AACTCATCTCAGAAGAGGATCTGAATAGCGCCGTCGACCATCATCATCAT<br>CATCATTGAGTTTAAACGGTCTCCAGCTTGGCTGTTTTGGCGGATGAGA<br>GAAGATTTTCAGCCTGATACAGATTAAATCAGAACGCAGAAGCGGTCTG<br>ATAAAACAGAATTTGCCTGGCGGCAGTAGCGCGGTGGTCCCACCTGACC<br>CCATGCCGAACTCAGAAGTGAAACGCCGTAGCGCCGATGGTAGTGTGG<br>GGTCTCCCCATGCGAGAGTAGGGAACTGCCAGGCATCAAATAAAACGAA<br>AGGCTCAGTCGAAAGACTGGGCCTTTCGTTTTATCTGTTGTTTGTCGGTG<br>AACTGGATCCCCAATTATTGAAGGCCGCTAACGCGGCCTTTTTTTGTTTC<br>TGGTCTGCCTTAATCAATGACTAAGAATTCGCGGCCGCTTCTAGAGCGT<br>CTCACTTCGGATCCGCTGCAGGCTTCTCAAGTCAAAAGCCTCCGGTCGG<br>AGGTTTTTGACTTTCTGCTATGGAGGTCAGGTATGATTTTTATGACAACTT<br>GACGGCTACATCATTCACTTTTTCTTCACAACCGGCACGGAACTCGCTC<br>GGGCTGGCCCCGGTGCATTTTTTAAATACCCGCGAGAAATAGAGTTGAT<br>CGTCAAAACCAACATTGCGACCGACGGTGGCGATAGGCATCCGGGTGG<br>TGCTCAAAAGCAGCTTCGCCTGGCTGATACGTTGGTCCTCGCGCCAGCT<br>TAAGACGCTAATCCCTAACTGCTGGCGGAAAAGATGTGACAGACGCGAC<br>GGCGACAAGCAAACATGCTGTGCGACGCTGGCGATATCAAAATTGCTGT<br>CTGCCAGGTGATCGCTGATGTACTGACAAGCCTCGCGTACCCGATTATC<br>CATCGGTGGATGGAGCGACTCGTTAATCGCTTCCATGCGCCGCAGTAAC<br>AATTGCTCAAGCAGATTTATCGCCAGCAGCTCCGAATAGCGCCCTTCCC<br>CTTGCCCGGCGTTAATGATTTGCCCAAACAGGTCGCTGAAATGCGGCTG<br>GTGCGCTTCATCCGGGCGAAAGAACCCCGTATTGGCAAATATTGACGGC |

CAGTTAAGCCATTCATGCCAGTAGGCGCGCGGACGAAAGTAAACCCACT
GGTGATACCATTCGCGAGCCTCCGGATGACGACCGTAGTGATGAATCTC
TCCTGGCGGGAACAGCAAAATATCACCCGGTCGGCAAACAAATTCTCGT
CCCTGATTTTTCACCACCCCCTGACCGCGAATGGTGAGATTGAGAATATA
ACCTTTCATTCCCAGCGGTCGGTCGATAAAAAAATCGAGATAACCGTTG
GCCTCAATCGGCGTTAAACCCGCCACCAGATGGGCATTAAACGAGTATC
CCGGCAGCAGGGGATCATTTTGCGCTTCAGCCATACTTTTCATACTCCC
GCCATTCAGAGAAGAAACCAATTGTCCATATTGCATCAGACATTGCCGTC
ACTGCGTCTTTTACTGGCTCTTCTCGCTAACCAAACCGGTAACCCCGCTT
ATTAAAAGCATTCTGTAACAAAGCGGGACCAAAGCCATGACAAAAACGC
GTAACAAAAGTGTCTATAATCACGGCAGAAAAGTCCACATTGATTATTTG
CACGGCGTCACACTTTGCTATGCCATAACATTTTTATCCATAAGATTAGC
GGATCCTACCTGACGCTTTTTATCGCAACTCTCTACTGTTTCTCCATACC
CGTTTTTTTGGGCTAGCCGACCTGTGGACGGATAAAGTCAGTTGTGAAA
TAAACGCGGCCATAGGCCGCGTAGTTCACTGCCGTATAGGCAGCTAAGA
AAAATGTGAACTGTATACATTCCCCGCAAGATAAAGGTGTAGAAATGTGA
GAGCGAAATGAGATGAGCGTTTTGTGAAATAAACGCGGCCATAGGCCGC
GTAGTTCACTGCCGTATAGGCAGCTAAGAAAAATGTGAACTGTATACATT
CCCCGCAAGATAAAGGTGTAGAAATGTGAGAGCGAAATGAGATGAGTTC
CTTGTGAAATAAACGCGGCCATAGGCCGCGTAGTTCACTGCCGTATAGG
CAGCTAAGAAAAATGTGAACTGTATACATTCCCCGCAAGATAAAGGTGTA
GAAATGTGAGAGCGAAATGAGATGAGTCATTTGTGAAATAAACGCGGCC
ATAGGCCGCGTAGTTCACTGCCGTATAGGCAGCTAAGAAAAATGTGAAC
TGTATACATTCCCCGCAAGATAAAGGTGTAGAAATGTGAGAGCGAAATG
AGATGAGAATGTTGTGAAATAAACGCGGCCATAGGCCGCGTAGTTCACT
GCCGTATAGGCAGCTAAGAAAAATGTGAACTGTATACATTCCCCGCAAG
ATAAAGGTGTAGAAATGTGAGAGCGAAATGAGATGAGCGTTTTGTGAAAT
AAACGCGGCCATAGGCCGCGTAGTTCACTGCCGTATAGGCAGCTAAGAA
AAATGTGAACTGTATACATTCCCCGCAAGATAAAGGTGTAGAAATGTGAG
AGCGAAATGAGATGAGTTCCTTGTGAAATAAACGCGGCCATAGGCCGCG
TAGTTCACTGCCGTATAGGCAGCTAAGAAAAATGTGAACTGTATACATTC
CCCGCAAGATAAAGGTGTAGAAATGTGAGAGCGAAATGAGATGAGTCAT
TTGTGAAATAAACGCGGCCATAGGCCGCGTAGTTCACTGCCGTATAGGC
AGCTAAGAAAAATGTGAACTGTATACATTCCCCGCAAGATAAAGGTGTAG
AAATGTGAGAGCGAAATGAGATGAGTGGCCAAAGCCCGCCGAAAGGCG
GGCTTTTTTTTGCTGTTCCTGAGACGTACTAGTAGGCCGCTGCAGTTAAT
CACTGATTAACTCGGTACCAAATTCCAGAAAAGAGGCCTCCCGAAAGGG
GGGCCTTTTTTCGTTTTGGTCCCTCGAGTCTAGACTGCAGTTGATCGGC
GTTAATATTTTGTTAAAATTCGCGTTAAATTTTTGTTAAATCAGCTCATTTT
TTAACCAATAGGCCGACTGCGATGAGTGGCAGGGCGGGGCGTAATTTTT
TTAAGGCAGTTATTGGTGCCCTTAAACGCGTTCTTTACAGCTAGCTCAGT
CCTAGGTATTATGCTAGCAGATCTAAAGAGGAGAAAGGATCTATGGACC
ACTACCTCGACATTCGCTTGCGACCGGACCCGGAATTTCCCCCGGCGCA

ACTCATGAGCGTGCTCTTCGGCAAGCTCCACCAGGCCCTGGTGGCACA
GGGCGGGGACAGGATCGGCGTGAGCTTCCCCGACCTCGACGAAAGCC
GCTCCCGGCTGGGCGAGCGCCTGCGCATTCATGCCTCGGCGGACGAC
CTTCGTGCCCTGCTCGCCCGGCCCTGGCTGGAAGGGTTGCGGGACCAT
CTGCAATTCGGAGAACCGGCAGTCGTGCCTCACCCCACACCGTACCGT
CAGGTCAGTCGGGTTCAGGCGAAAAGCAATCCGGAACGCCTGCGGCGG
CGGCTCATGCGCCGGCACGATCTGAGTGAGGAGGAGGCTCGGAAACGC
ATTCCCGATACGGTCGCGAGAGCCTTGGACCTGCCCTTCGTCACGCTAC
GCAGCCAGAGCACCGGACAGCACTTCCGTCTCTTCATCCGCCACGGGC
CGTTGCAGGTGACGGCAGAGGAAGGAGGATTCACCTGTTACGGGTTGA
GCAAAGGAGGTTTCGTTCCCTGGTTCTGATAACTCGAGTAAGGATCTCC
AGGCATCAAATAAAACGAAAGGCTCAGTCGAAAGACTGGGCCTTTCGTT
TTATCTGTTGTTTGTCGGTGAACTTACTCGAGTCTAGACTGCAGCTGAAA
CTGGTGCTACGCCTGAATAAGTGATAATAAGCGGATGAATGGCAGAAAT
TCGAAAGCAAATTCGACCCGGTCGTCGGTTCAGGGCAGGGTCGTTAAAT
AGCCGCTTATGTCTATTGCTGGTTTACCGGTTTATTGACTACCGGAAGCA
GTGTGACCGTGTGCTTCTCAAATGCCTGAGGCCAGTTTGCTCAGGCTCT
CCCCGTGGAGGTAATAATTGACGATATGATCATTTATTCTGCCTCCCAGA
GCCTGATAAAAACGGTGAATCCGTTAGCGAGGTGCCGCCGGCTTCCATT
CAGGTCGAGGTGGCCCGGCTCCATGCACCGCGACGCAACGCGGGGAG
GCAGACAAGGTATAGGGCGGCGAGGCGGCTACAGCCGATAGTCTGGAA
CAGCGCACTTACGGGTTGCTGCGCAACCCAAGTGCTACCGGCGCGGCA
GCGTGACCCGTGTCGGCGGCTCCAACGGCTCGCCATCGTCCAGAAAAC
ACGGCTCATCGGGCATCGGCAGGCGCTGCTGCCCGCGCCGTTCCCATT
CCTCCGTTTCGGTCAAGGCTGGCAGGTCTGGTTCCATGCCCGGAATGC
CGGGCTGGCTGGGCGGCTCCTCGCCGGGGCCGGTCGGTAGTTGCTGC
TCGCCCGGATACAGGGTCGGGATGCGGCGCAGGTCGCCATGCCCCAAC
AGCGATTCGTCCTGGTCGTCGTGATCAACCACCACGGCGGCACTGAACA
CCGACAGGCGCAACTGGTCGCGGGGCTGGCCCCACGCCACGCGGTCA
TTGACCACGTAGGCCGACACGGTGCCGGGGCCGTTGAGCTTCACGACG
GAGATCCAGCGCTCGGCCACCAAGTCCTTGACTGCGTATTGGACCGTCC
GCAAAGAACGTCCGATGAGCTTGGAAAGTGTCTTCTGGCTGACCACCAC
GGCGTTCTGGTGGCCCATCTGCGCCACGAGGTGATGCAGCAGCATTGC
CGCCGTGGGTTTCCTCGCAATAAGCCCGGCCCACGCCTCATGCGCTTT
GCGTTCCGTTTGCACCCAGTGACCGGGCTTGTTCTTGGCTTGAATGCCG
ATTTCTCTGGACTGCGTGGCCATGCTTATCTCCATGCGGTAGGGTGCCG
CACGGTTGCGGCACCATGCGCAATCAGCTGCAACTTTTCGGCAGCGCG
ACAACAATTATGCGTTGCGTAAAAGTGGCAGTCAATTACAGATTTTCTTTA
ACCTACGCAATGAGCTATTGCGGGGGGTGCCGCAATGAGCTGTTGCGT
ACCCCCCTTTTTTAAGTTGTTGATTTTTAAGTCTTTCGCATTTCGCCCTAT
ATCTAGTTCTTTGGTGCCCAAAGAAGGGCACCCCTGCGGGGTTCCCCCA
CGCCTTCGGCGCGGCTCCCCCTCCGGCAAAAAGTGGCCCCTCCGGGG
CTTGTTGATCGACTGCGCGGCCTTCGGCCTTGCCCAAGGTGGCGCTGC

CCCCTTGGAACCCCCGCACTCGCCGCCGTGAGGCTCGGGGGGCAGGC
GGGCGGGCTTCGCCTTCGACTGCCCCCACTCGCATAGGCTTGGGTCGT
TCCAGGCGCGTCAAGGCCAAGCCGCTGCGCGGTCGCTGCGCGAGCCTT
GACCCGCCTTCCACTTGGTGTCCAACCGGCAAGCGAAGCGCGCAGGCC
GCAGGCCGGAGGCTTTTCCCCAGAGAAAATTAAAAAAATTGATGGGGCA
AGGCCGCAGGCCGCGCAGTTGGAGCCGGTGGGTATGTGGTCGAAGGC
TGGGTAGCCGGTGGGCAATCCCTGTGGTCAAGCTCGTGGGCAGGCGCA
GCCTGTCCATCAGCTTGTCCAGCAGGGTTGTCCACGGGCCGAGCGAAG
CGAGCCAGCCGGTGGCCGCTCGCGGCCATCGTCCACATATCCACGGGC
TGGCAAGGGAGCGCAGCGACCGCGCAGGGCGAAGCCCGGAGAGCAAG
CCCGTAGGGCGCCGCAGCCGCCGTAGGCGGTCACGACTTTGCGAAGCA
AAGTCTAGTGAGTATACTCAAGCATTGAGTGGCCCGCCGGAGGCACCG
CCTTGCGCTGCCCCCGTCGAGCCGGTTGGACACCAAAAGGGAGGGGCA
GGCATGGCGGCATACGCGATCATGCGATGCAAGAAGCTGGCGAAAATG
GGCAACGTGGCGGCCAGTCTCAAGCACGCCTACCGCGAGCGCGAGAC
GCCCAACGCTGACGCCAGCAGGACGCCAGAGAACGAGCACTGGGCGG
CCAGCAGCACCGATGAAGCGATGGGCCGACTGCGCGAGTTGCTGCCAG
AGAAGCGGCGCAAGGACGCTGTGTTGGCGGTCGAGTACGTCATGACGG
CCAGCCCGGAATGGTGGAAGTCGGCCAGCCAAGAACAGCAGGCGGCG
TTCTTCGAGAAGGCGCACAAGTGGCTGGCGGACAAGTACGGGGCGGAT
CGCATCGTGACGGCCAGCATCCACCGTGACGAAACCAGCCCGCACATG
ACCGCGTTCGTGGTGCCGCTGACGCAGGACGGCAGGCTGTCGGCCAA
GGAGTTCATCGGCAACAAAGCGCAGATGACCCGCGACCAGACCACGTT
TGCGGCCGCTGTGGCCGATCTAGGGCTGCAACGGGGCATCGAGGGCA
GCAAGGCACGTCACACGCGCATTCAGGCGTTCTACGAGGCCCTGGAGC
GGCCACCAGTGGGCCACGTCACCATCAGCCCGCAAGCGGTCGAGCCAC
GCGCCTATGCACCGCAGGGATTGGCCGAAAAGCTGGGAATCTCAAAGC
GCGTTGAGACGCCGGAAGCCGTGGCCGACCGGCTGACAAAAGCGGTTC
GGCAGGGGTATGAGCCTGCCCTACAGGCCGCCGCAGGAGCGCGTGAG
ATGCGCAAGAAGGCCGATCAAGCCCAAGAGACGGCCCGAGACCTTCGG
GAGCGCCTGAAGCCCGTTCTGGACGCCCTGGGGCCGTTGAATCGGGAT
ATGCAGGCCAAGGCCGCCGCGATCATCAAGGCCGTGGGCGAAAAGCTG
CTGACGGAACAGCGGGAAGTCCAGCGCCAGAAACAGGCCCAGCGCCA
GCAGGAACGCGGGCGCGCACATTTCCCCGAAAAGTGCCACCTGGGATG
AATGTCAGCTACTGGGCTATCTGGACAAGGGAAAACGCAAGCGCAAAGA
GAAAGCAGGTAGCTTGCAGTGGGCTTACATGGCGATAGCTAGACTGGG
CGGTTTTATGGACAGCAAGCGAACCGGAATTGCCAGCTGGGGCGCCCT
CTGGTAAGGTTGGGAAGCCCTGCAAAGTAAACTGGATGGCTTTCTTGCC
GCCAAGGATCTGATGGCGCAGGGGATCAAGATCTGATCAAGAGACAGG
ATGAGGATCGTTTCGCATGATTGAACAAGATGGATTGCACGCAGGTTCT
CCGGCCGCTTGGGTGGAGAGGCTATTCGGCTATGACTGGGCACAACAG
ACAATCGGCTGCTCTGATGCCGCCGTGTTCCGGCTGTCAGCGCAGGGG
CGCCCGGTTCTTTTTGTCAAGACCGACCTGTCCGGTGCCCTGAATGAAC

TGCAGGACGAGGCAGCGCGGCTATCGTGGCTGGCCACGACGGGCGTT
CCTTGCGCAGCTGTGCTCGACGTTGTCACTGAAGCGGGAAGGGACTGG
CTGCTATTGGGCGAAGTGCCGGGGCAGGATCTCCTGTCATCTCACCTTG
CTCCTGCCGAGAAAGTATCCATCATGGCTGATGCAATGCGGCGGCTGCA
TACGCTTGATCCGGCTACCTGCCCATTCGACCACCAAGCGAAACATCGC
ATCGAGCGAGCACGTACTCGGATGGAAGCCGGTCTTGTCGATCAGGAT
GATCTGGACGAAGAGCATCAGGGGCTCGCGCCAGCCGAACTGTTCGCC
AGGCTCAAGGCGCGCATGCCCGACGGCGAGGATCTCGTCGTGACCCAT
GGCGATGCCTGCTTGCCGAATATCATGGTGGAAAATGGCCGCTTTTCTG
GATTCATCGACTGTGGCCGGCTGGGTGTGGCGGACCGCTATCAGGACA
TAGCGTTGGCTACCCGTGATATTGCTGAAGAGCTTGGCGGCGAATGGG
CTGACCGCTTCCTCGTGCTTTACGGTATCGCCGCTCCCGATTCGCAGCG
CATCGCCTTCTATCGCCTTCTTGACGAGTTCTTCTGAGCGGGACTCTGG
GGTTCGAAATGACCGACCAAGCGACGCCCAACCTGCCATCACGAGATTT
CGATTCCACCGCCGCCTTCTATGAAAGGTTGGGCTTCGGAATCGTTTTC
CGGGACGCCGGCTGGATGATCCTCCAGCGCGGGGATCTCATGCTGGAG
TTCTTCGCCCACCCCCATGGGCAAATATTATACGCAAGGCGACAAGGTG
CTGATGCCGCTGGCGATTCAGGTTCATCATGCCGTTTGTGATGGCTTCC
ATGTCGGCAGAATGCTTAATGAATTACAACAGTTTTTATGCATGCGCCCA
ATACGCAAACCGCCTCTCCCCGCGCGTTGGCCGATTCATTAATGCAGCT
GGCACGACAGGTTTCCCGACTGGAAAGCGGGCAGTGAGCGCAACGCAA
TTAATGTGAGTTAGCTCACTCATTAGGCACCCCAGGCCTAGATTTCAGTG
CAATTTATCTCTTCAAATGTAGCACCTGAAGTCAGCCCCATACGATATAA
GTTGTAATTCTCATGTTTGACAGCTTATCATCGATAAGCTTCCGATGGCG
CGCCGAGAGGCTTTACACTTTATGCTTCCGGCT

**Supplementary Table 3. SP6 promoter sequences used in this study.**

| Promoter | Sequence (5' to 3') |
|---|---|
| pSP6_wt | ATTTAGGTGACACTATAGGG |
| pSP6_90 | ATTTAGGTGCCACTATAGGG |
| pSP6_50 | ATTTAGGTGACACTATGGGG |
| pSP6_20 | ATTTAGGTGACATTATAGGG |

**Supplementary Table 4. Golden Gate assembly primers and overhangs.** An example of the 8-copy RNA array assembly. BsmBI site: GAGACG, BsaI site: GGTCTC, Overhang: ctcg. Table corresponds to the naming conventions shown in Supplementary Figure 2. We note that some of the parts indicated in the table below (e.g., spacer, connectors) are not included in Supplementary Figure 2 for simplicity. The top section of the table indicates how to construct a 4-copy STAR array modular cloning (MoClo) part for position 3b. Position 4 can be used to create another 4-copy STAR array MoClo part using the same overhang logic indicated in the top section of the table. If constructing a multiplex array (Figure 4D) the STAR identity in positions 3b and 4 can be varied.

| **Assembly parts** | **Sequence (5' to 3')** |
|---|---|
| **Level 1 => Level 2 (Position 3b)** | |
| Backbone | - CGTCTCaaGAGACC - backbone - GGTCTCgtGAGACG - GFP dropout - |
| PCR products 1 | gtCGTCTCactcgtcag - Insulator+STAR - cgtttGAGACGtg |
| PCR products 2 | gtCGTCTCacgtt - Insulator+STAR - ttcctGAGACGtg |
| PCR products 3 | gtCGTCTCattcc - Insulator+STAR - tcattGAGACGtg |
| PCR products 4 | gtCGTCTCatcat - Insulator+STAR - aatgagagtGAGACGtg |
| **Level 2 => Level 3** | |
| Position 1 | GGTCTCgacgg - Connector* - cttcaGAGACC |
| Position 2 | GGTCTCgcttc - Promoter - cgcaaGAGACC |
| Position 3a | GGTCTCgcgca - Spacer/Target - tcagaGAGACC |
| Position 3b | GGTCTCgtcag - Inserts - aatgaGAGACC |
| Position 4 | GGTCTCgaatg - Inserts - tggcaGAGACC |
| Position 5 | GGTCTCgtggc - Terminator - gctgaGAGACC |
| Position 6 | GGTCTCggctg - Connector - tacaaGAGACC |
| Position 7 | GGTCTCgtaca - Resistance & Origin of replication - acggaGAGACC |

*Connectors contain BsmBI sites allowing further assembly

**Supplementary Table 5. Strains, growth conditions, and electroporation settings used in this study.**

| Species | Strain name | Culture media | Culture temperature (ºC) | Kanamycin conc. (µg/mL) | Electroporation setting (kV) |
|---|---|---|---|---|---|
| *Escherichia coli* | NEB Turbo (cloning strain) | LB | 37 | 100 | - |
| *Escherichia coli* | TG1 | LB | 37 | 100 | - |
| *Shewanella oneidensis* | MR-1 | LB | 30 | 50 | 1.2 |
| *Pseudomonas fluorescens* | A506 | LB | 30 | 50 | 2.5 |
| *Pseudomonas putida* | F1 | LB | 30 | 50 | 1.25 |
| *Pseudomonas stutzeri* | JM300 | LB | 30 | 50 | 2.5 |
| *Vibrio natriegens* | Vmax | LB3 | 37 | 200 | 0.7 |

**Supplementary Table 6. Regulatory RNA array insulator sequences.**

| Insulator | Sequence (5' to 3') |
|---|---|
| PlmJ | AGUCAUAAGUCUGGGCUAAGCCCACUGAUGAGUCGCUGAAAUGCGACGAAACUUAUGA |
| csy4hp | GUUCACUGCCGUAUAGGCAGCUAAGAAA |
| shcsy4hp | UUGUGAAAUAAACGCGGCCAUAGGCCGCGUAGUUCACUGCCGUAUAGGCAGCUAAGAAA |
| PlmJ-SATR50x4 (Example sequence used for NUPACK simulation) | AGUCAUAAGUCUGGGCUAAGCCCACUGAUGAGUCGCUGAAAUGCGACGAAACUUAUGAUGAACUGUAUACAUUCCCCGCAAAGUGCCUAUCUGUCGUCGUGUUAUCUUUAUGUUUCUGGCGUUAGUCAUAAGUCUGGGCUAAGCCCACUGAUGAGUCGCUGAAAUGCGACGAAACUUAUGAUGAACUGUAUACAUUCCCCGCAAAGUGCCUAUCUGUCGUCGUGUUAUCUUUAUGUUUCUGGUUCCAGUCAUAAGUCUGGGCUAAGCCCACUGAUGAGUCGCUGAAAUGCGACGAAACUUAUGAUGAACUGUAUACAUUCCCCGCAAAGUGCCUAUCUGUCGUCGUGUUAUCUUUAUGUUUCUGGUCAUAGUCAUAAGUCUGGGCUAAGCCCACUGAUGAGUCGCUGAAAUGCGACGAAACUUAUGAUGAACUGUAUACAUUCCCCGCAAAGUGCCUAUCUGUCGUCGUGUUAUCUUUAUGUUUCUGG |

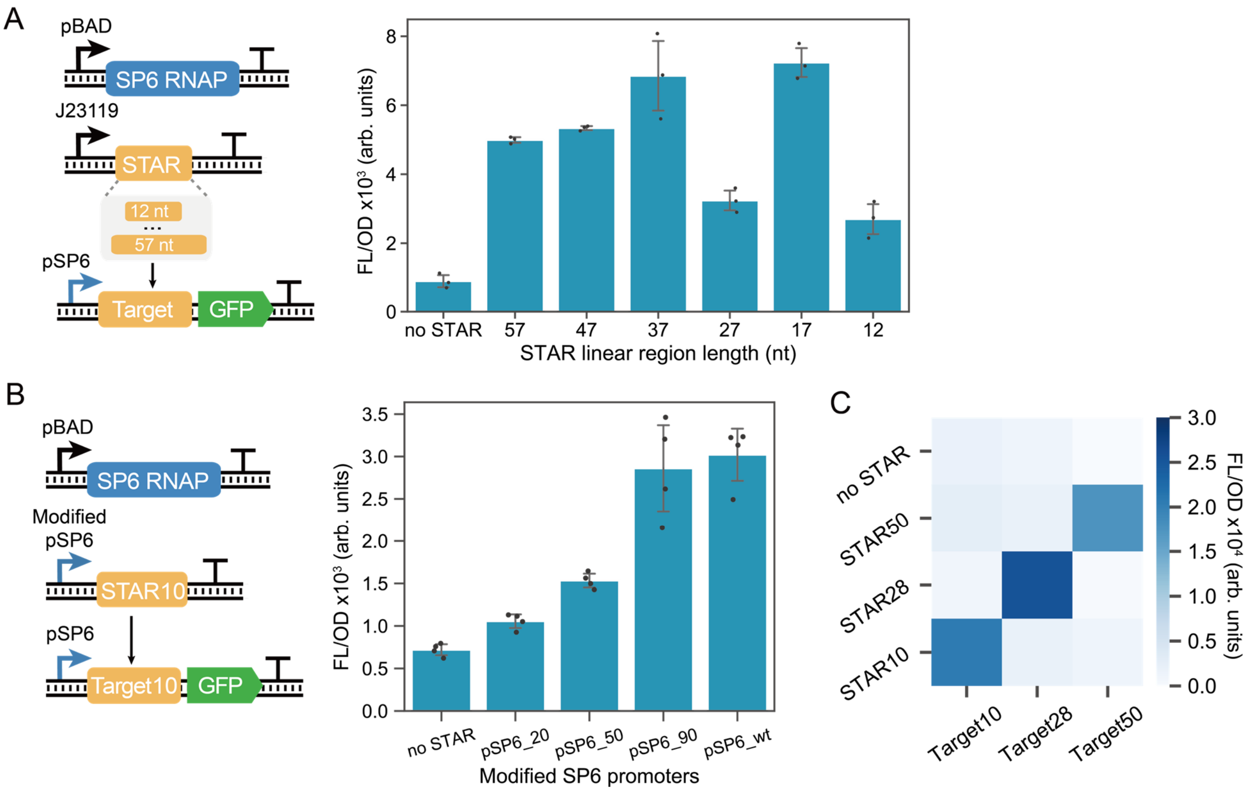


**Supplementary Fig. 1. Characterization of a STAR system that uses the SP6 RNA polymerase.** (A) Investigating the impact of length on the SP6 STAR system. STAR systems with different length linear regions (12 to 57 nucleotides [nt]) are used to activate a target RNA-controlled GFP that is transcribed using an SP6 promoter. The left panel shows a schematic of genetic circuitry and the right panel shows fluorescent characterization in *E. coli* cells transformed with corresponding plasmids. The results show a length of ~40 nt to be optimal which is comparable to prior STAR characterization results using the native *E. coli* RNAP[1]. Bars show mean values and error bars represent s.d. of $n = 3$ biological replicates shown as points. (B) Investigating the tunability of SP6-controlled STARs. STAR systems transcribed using only SP6 RNAP. The left panel shows a schematic of genetic circuitry and the right panel shows fluorescent characterization in *E. coli* cells transformed with corresponding plasmids. Variable strength SP6 promoters[2] are used to produce the STAR and a constant SP6 promoter is used to produce the target RNA. Bars show mean values and error bars represent s.d. of $n = 4$ biological replicates shown as points. (C) Investigating the orthogonality of SP6 STARs. Fluorescent characterization of cells transformed with different combinations of plasmids encoding cognate and non-cognate STAR and target RNA pairs. Cognate pairs are across the diagonal and the no STAR control is in the top row. Heatmap shows the mean values of $n = 4$ biological replicates.

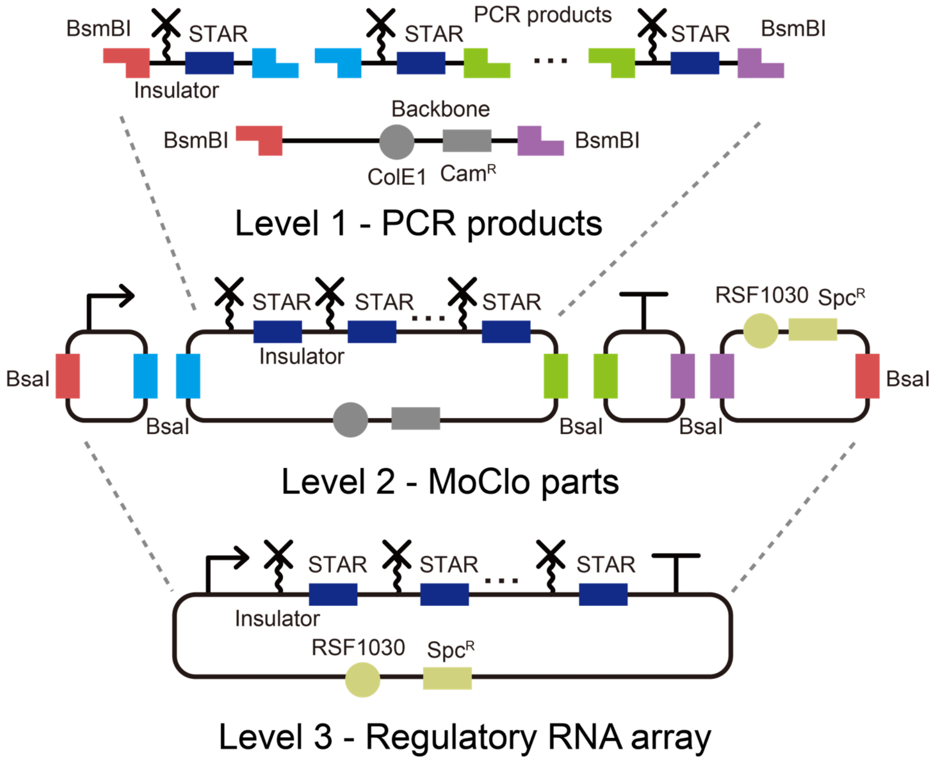


**Supplementary Fig. 2. A modular cloning (MoClo) approach for regulatory RNA arrays.** The regulatory RNA array is constructed through a 3-level modular cloning approach. PCR products are made with primers containing the recognition site for type 2 restriction enzyme BsmBI. The copy number of STARs can be altered by adjusting the number of PCR fragments with different overhangs. The PCR products are then assembled as MoClo part plasmids containing BsaI sites for long-term storage. Next, the MoClo parts are further assembled as the target plasmid with full regulatory RNA array cassette. We note that some of the parts indicated in Supplementary Table 4 (*e.g.,* spacer, connectors) are not included in this for simplicity.

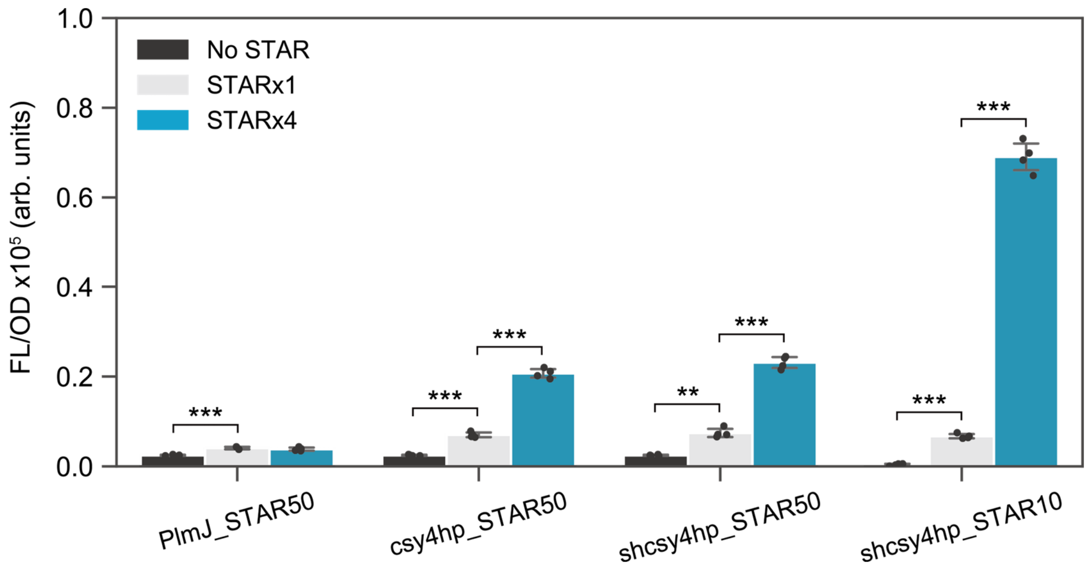


**Supplementary Fig. 3. Insulator screening for regulatory RNA arrays.** Fluorescent characterization of regulatory RNA arrays using PlmJ, csy4 hairpin, strong-hairpin csy4 hairpin as insulators. STAR50 and STAR10 arrays are tested with either x1, x4 or no copies. Bars show mean values and error bars represent s.d. of $n = 4$ biological replicates shown as points. Two-tailed t-tests assuming unequal variance were used and the significance are marked by asterisks indicating $p<.05$ (*), $p<.01$ (**), $p<.001$ (***).

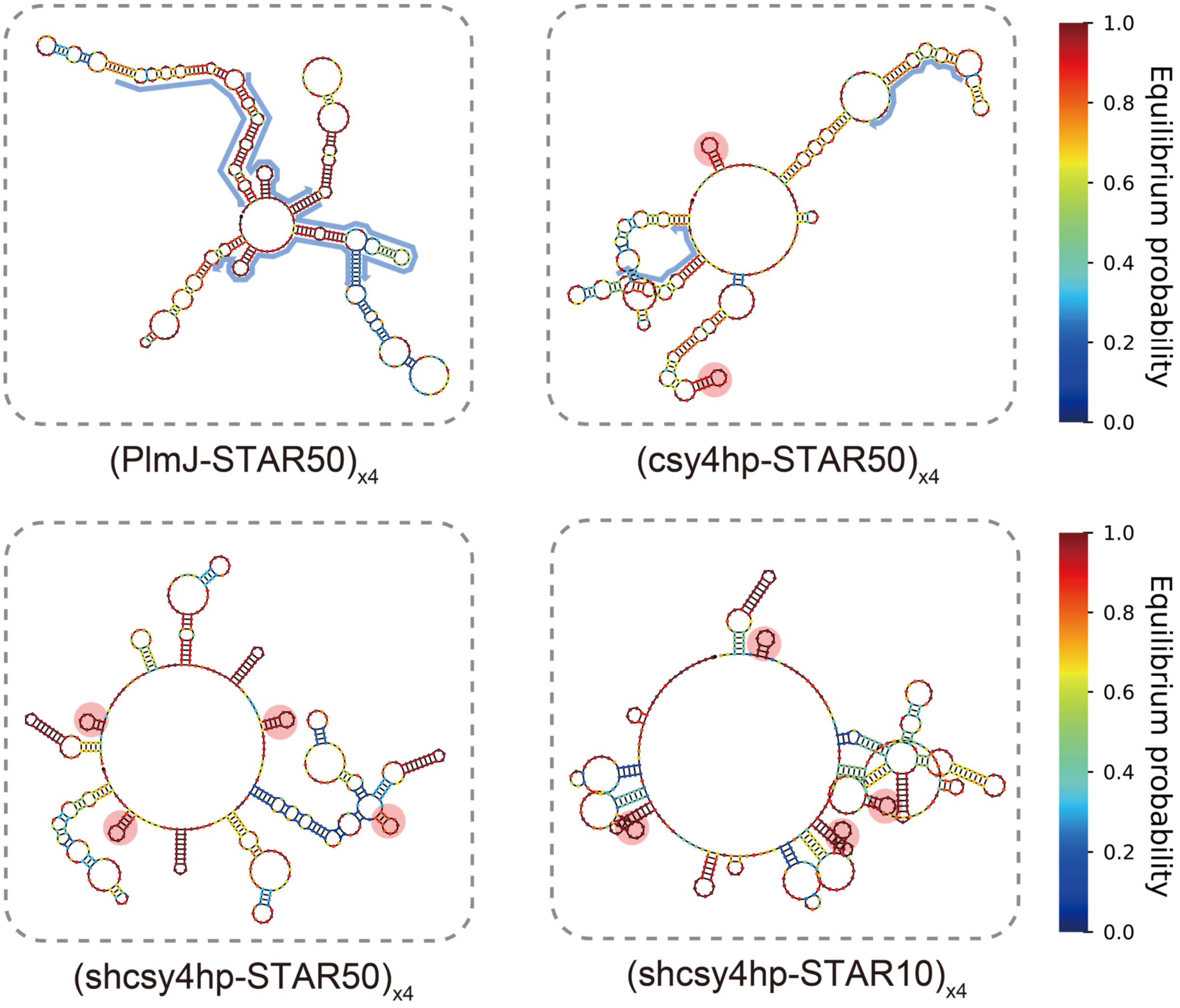


**Supplementary Fig. 4. NUPACK analysis of regulatory RNA arrays with different insulators.** Secondary structure predictions of 4-copy STAR arrays insulated with ribozyme PlmJ, ribonuclease site csy4 (csy4hp), and a strong-hairpin csy4 (shcsy4hp). Predictions use the online version of NUPACK[3]. The color of each nucleotide represents the equilibrium probability for each nucleotide to be in that state (e.g., paired/unpaired). Correctly folded insulator structures are labeled with red shading and insulator sequences that do not form the expected structure are indicated with blue shading.

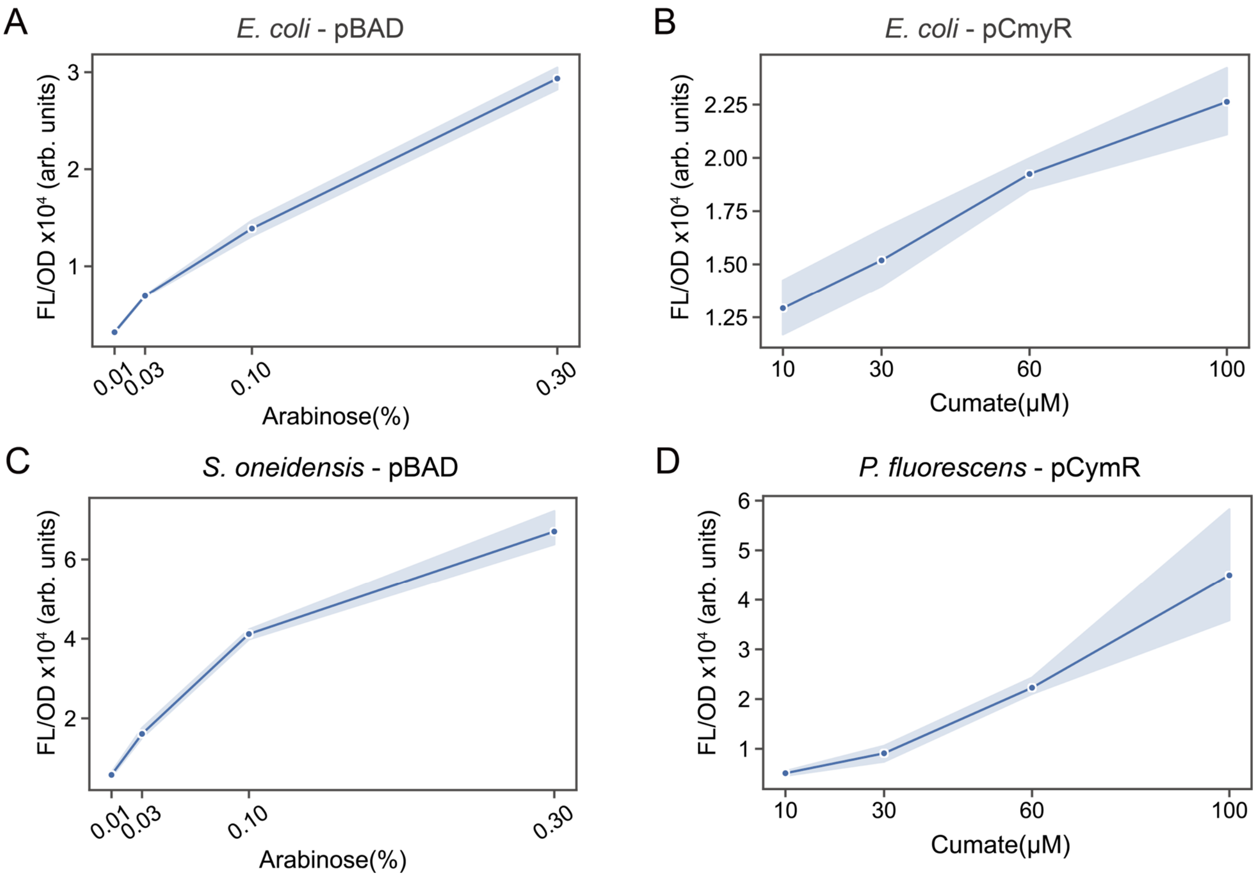


**Supplementary Fig. 5. Calibration curves of inducible promoters to relate promoter strength to transcriptional output in different bacterial species.** Fluorescent characterization of cells transformed with plasmids encoding a GFP driven by either pCymR or pBAD under different concentrations of inducer. Graphs show data from (A) pBAD in *E. coli*, (B) pCymR in *E. coli,* (C) pBAD in *S. oneidensis,* and (D) pCmyR in *P. fluorescens*. Each point shows the mean value and the shade represents s.d. of $n = 4$ biological replicates.

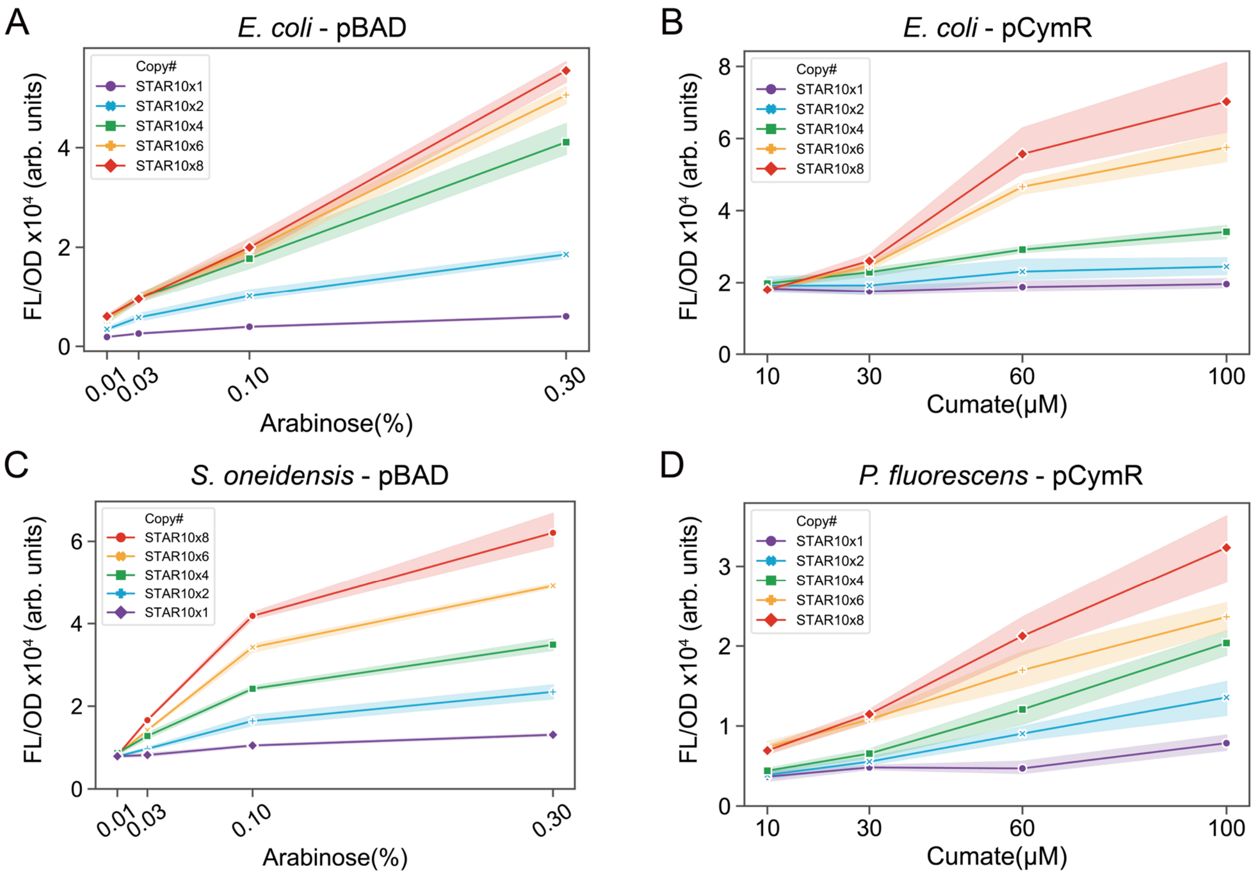


**Supplementary Fig. 6. The induction curves of the regulatory RNA arrays before the promoter activity normalization.** Fluorescent characterization of cells transformed with plasmids encoding regulatory RNA arrays with 1, 2, 4, 6, and 8 copies of STARs driven by either pCymR or pBAD under different concentrations of inducer. Graphs show data from (A) pBAD in *E. coli*, (B) pCymR in *E. coli,* (C) pBAD in *S. oneidensis,* and (D) pCmyR in *P. fluorescens*. Each point shows the mean value and the shade represents s.d. of *n* = 4 biological replicates.

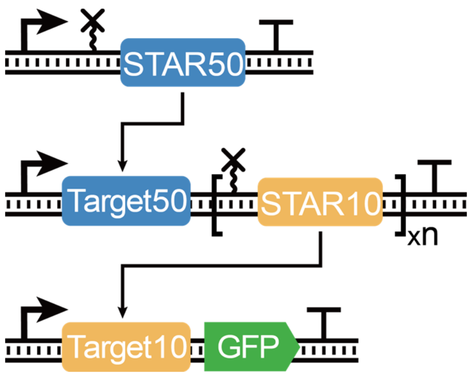


**Supplementary Fig. 7. The schematic of an RNA activation-activation cascade.** The RNA cascade is built with an orthogonal pair STAR10 and STAR50. STAR50 is constitutively transcribed and activates the transcription of the regulatory RNA array composed of variable copies of STAR10. STAR10 production leads to the activation of Target10, which in turn activates the production of GFP.

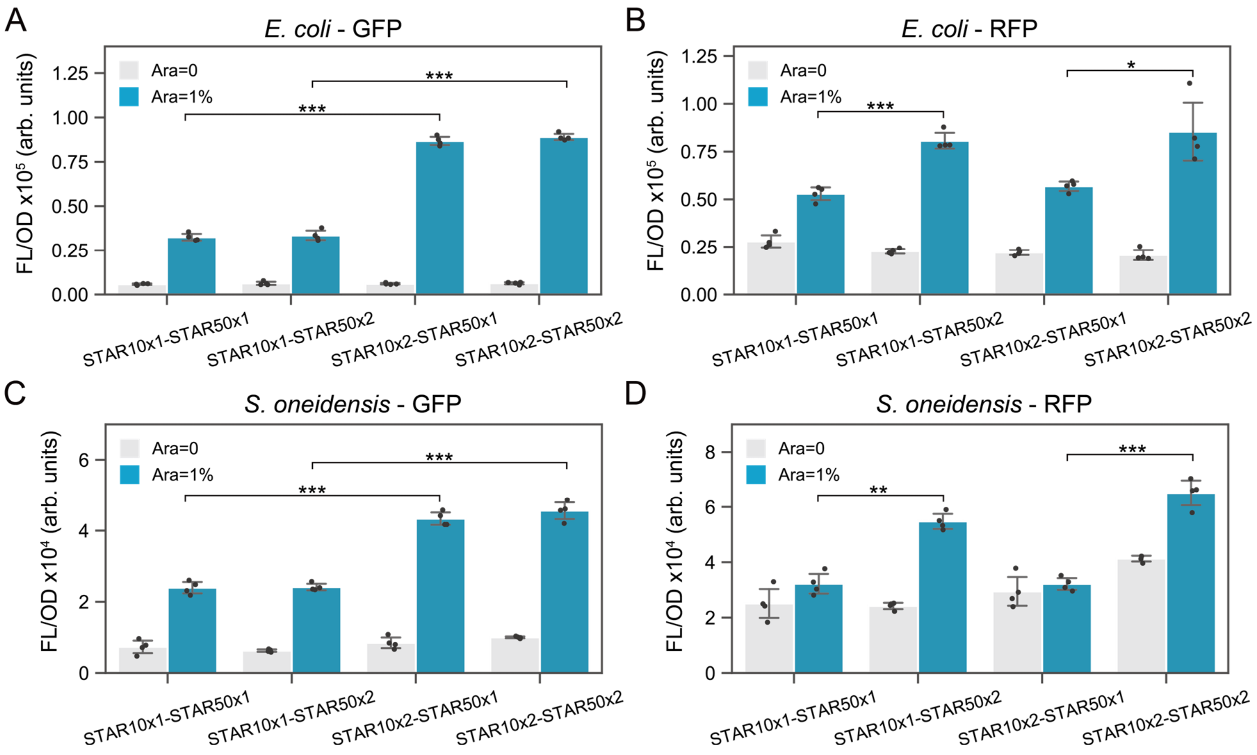


**Supplementary Fig. 8. Fluorescent characterization of multiplex RNA arrays.** Fluorescent characterization of the multiplex RNA arrays with the 2x2 matrix of 4 possible STAR10 and STAR50 combinations. The graphs show uninduced and induced (arabinose=1%) data from (A) GFP characterization in *E. coli*, (B) RFP characterization in *E. coli*, (C) GFP characterization in *S. oneidensis*, (D) RFP characterization in *S. oneidensis*. Bars show mean values and error bars represent s.d. of $n = 4$ biological replicates shown as points. Two-tailed t-tests assuming unequal variance were used and the significance are marked by asterisks indicating $p<.05$ (*), $p<.01$ (**), $p<.001$ (***).

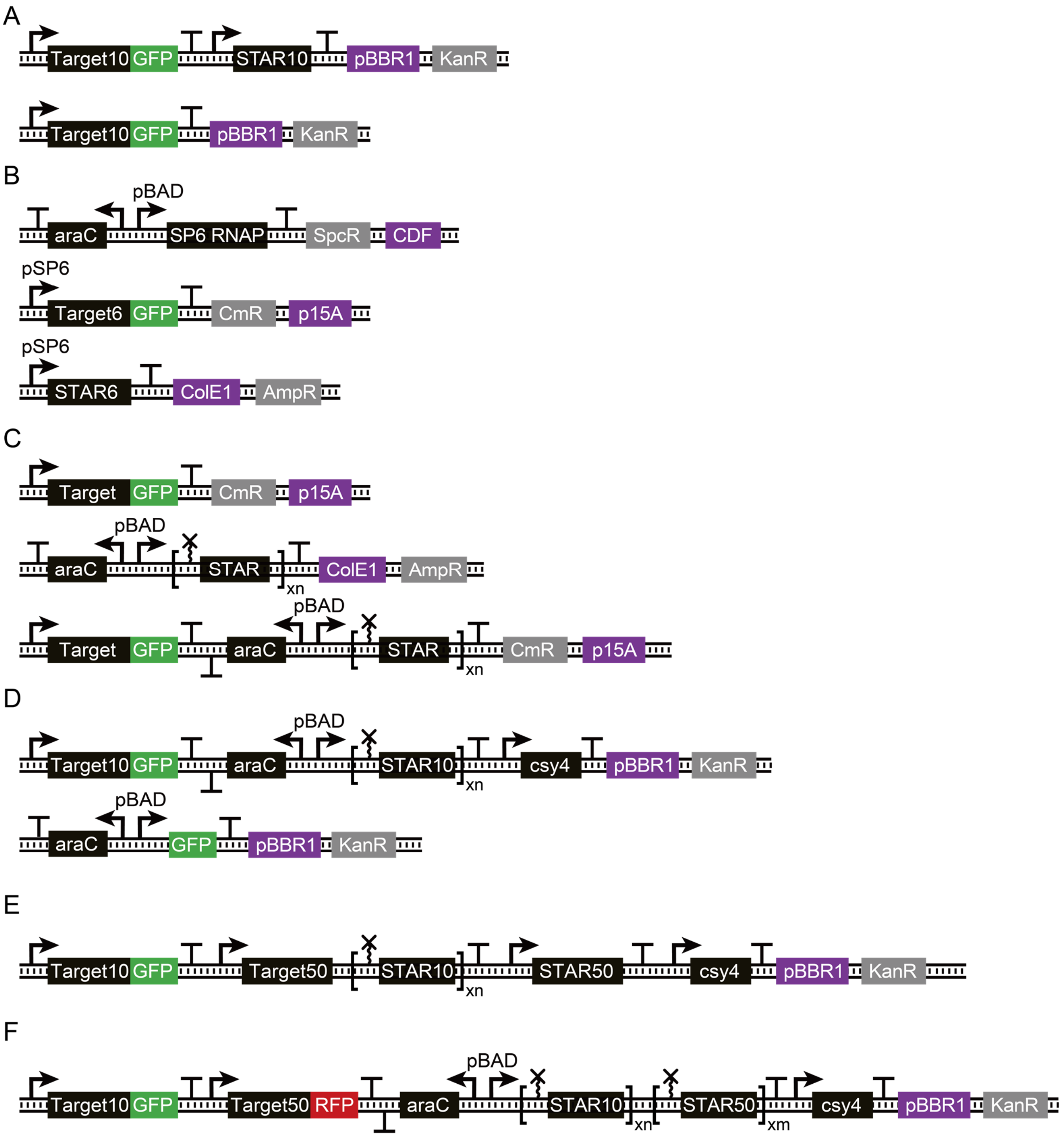


**Supplementary Fig. 9. The schematics of representative plasmids used in this study.** The example plasmid set used in (A) STAR for Gram-negative RNAP, (B) SP6 RNAP-driven STAR, (C) regulatory RNA array insulation screening, (D) regulatory RNA array and standard inducible GFP on broad host range plasmid, (E) single-plasmid RNA cascade, (F) multiplex regulatory RNA array.

**Supplementary Note 1. Mathematical model of regulatory RNA arrays.**
The model consists of two main process: (1) Csy4 processing of the RNA array and (2) the regulation of small transcription activating RNA (STAR) system. For the rest of this note, we will use capital letter to indicate chemical species, and lower case to denote the corresponding concentration. For example, species $X$ has concentration $x$.

**Modeling Csy4 cleaving.** First, we define $P^*$ to describe the unbound molecular specie Csy4, $P$ for bound Csy4 to RNA array, and $X_n$ for the full RNA array with $n$ repeats. We consider a production rate constant $\alpha$ and $\theta$ for $P^*$ and $X_n$ respectively. A degradation rate constant $\delta$ and $\phi$ for $P^*$ and $X_n$. $n$ unbound proteins $P^*$ can bind/unbinding to a single RNA array $X_n$ and form a complex $P$ with a rate constant $k_p^+$ and $k_p^-$. Finally, the state $P$ can cleave and produce $n$ copies of STAR $X$ at a rate constant $r_p$, which decays at a rate constant $\phi$. We summarize the reactions below:

| | | |
|---|---|---|
| $\emptyset \xrightarrow{\alpha} P^*$ | $P^* \xrightarrow{\delta} \emptyset$ | Production/Degradation |
| $\emptyset \xrightarrow{\theta} X_n$ | $X_n \xrightarrow{\phi} \emptyset$ | Production/Degradation |
| $X_n + nP^* \underset{\kappa_p^-}{\overset{\kappa_p^+}{\rightleftharpoons}} P$ | $P \xrightarrow{r_p} nX.$ | Binding/Unbinding/Cleaving |
| | $X \xrightarrow{\phi} \emptyset$ | Degradation |

As a result, we can write down the Ordinary Differential Equations (ODEs) by using the law of mass action:

$$\dot{p}^* = \alpha - \delta p^* - n * k_p^+ x_n p^* + n k_p^- p \quad (1)$$
$$\dot{x}_n = \theta - \phi x_n - k_p^+ x_n p^* + k_p^- p \quad (2)$$
$$\dot{p} = k_p^+ x_n p^* - k_p^- p - r_p p - \delta p \quad (3)$$
$$\dot{x} = n r_p p - \phi x \quad (4)$$

**Modeling STAR system.** A cleaved STAR $X$ can bind to the target RNA complex $C^*$ in which transcription is blocked and form a complex $C$ to activate transcription, with a binding and unbinding rate $k_c^+$, and $k_c^-$. These interactions will modify the model for $x$, equation (4). The activated complex $C$ then transcribes the output $Y$ at the rate of $r_c$. Also, we consider leakiness when the blocked transcription complex $C^*$ can produce an output species $Y$ at a rate constant $r_0$. Finally, abortion can be incorporated when the activate transcription complex $C$ become $C^*$ at a rate constant $\xi$. We summarize the chemical reactions below:

| | | |
|---|---|---|
| $X + C^* \underset{\kappa_c^-}{\overset{\kappa_c^+}{\rightleftharpoons}} C$ | $C \xrightarrow{r_c} C^* + Y.$ | Activation |
| $Y \xrightarrow{\phi} \emptyset$ | | Decay |
| $C^* \xrightarrow{r_0} Y$ | $C \xrightarrow{\xi} C^*$ | Leakiness/Abortion |

Following law mass of action, we can find the ODEs:

$$\dot{x} = n r_p p - \phi x - \kappa_c^+ x c^* + \kappa_c^- c \quad (5)$$
$$\dot{c} = \kappa_c^+ x c^* - \kappa_c^- c - r_c c - \xi c \quad (6)$$
$$\dot{y} = r_0 c^* + r_c c - \delta y \quad (7)$$

with a mass conservation of the total amount of DNA complex constant, $c + c^* = c^{tot}$.

**Numerical Simulations.** Below we list the parameters used in the model for simulations.

| Parameter | Value | Annotation | Other studies |
|---|---|---|---|
| $\phi, \delta$ (1/s) | $2.7 \times 10^{-4}$ | degradation | $10^{-4} - 10^{-3}$ [4] |
| $\theta$ (M/s) | $2.7 \times 10^{-10}$ | transcription | $2.8 \times 10^{-11} - 2.8 \times 10^{-8}$ [5,6] |
| $\alpha$ (M/s) | $5.4 \times 10^{-10}$ | transcription | $2.8 \times 10^{-11} - 2.8 \times 10^{-8}$ [5,6] |
| $k_p^+, k_c^+$ (/M/s) | $2.7 \times 10^{4}$ | binding | $10^4 - 10^6$ [7,8] |
| $k_p^-, k_c^-$ (1/s) | $2.7 \times 10^{-4}$ | unbinding | |
| $r_p, r_c$ (1/s) | $2.7 \times 10^{-3}$ | complex transcription | $0.05 - 0.2$ [9]* |
| $r_0$ (1/s) | $2.7 \times 10^{-4}$ | leakiness | |
| $c^{tot}$ (nM) | $1000$ | total DNA | |
| $\xi$ (1/s) | $0$ | abortion rate | |

* Estimated from the average pause-free velocity of RNAP at saturating concentrations of nucleotide triphosphates (NTP) for an RNA length of 200 nt.
